# Generating accurate 3D gaze vectors using synchronized eye tracking and motion capture

**DOI:** 10.1101/2021.10.22.465332

**Authors:** Scott A. Stone, Quinn A. Boser, T. Riley Dawson, Albert H. Vette, Jacqueline S. Hebert, Patrick M. Pilarski, Craig S. Chapman

## Abstract

Assessing gaze behaviour during real-world tasks is difficult; dynamic bodies moving through dynamic worlds make gaze analysis difficult. Current approaches involve laborious coding of pupil positions. In settings where motion capture and mobile eye tracking are used concurrently in naturalistic tasks, it is critical that data collection be simple, efficient, and systematic. One solution is to combine eye tracking with motion capture to generate 3D gaze vectors. When combined with tracked or known object locations, 3D gaze vector generation can be automated. Here we use combined eye and motion capture and explore how linear regression models generate accurate 3D gaze vectors. We compare spatial accuracy of models derived from four short calibration routines across three pupil data inputs: the efficacy of calibration routines were assessed, a validation task requiring short fixations on task-relevant locations, and a naturalistic object interaction task to bridge the gap between laboratory and “in the wild” studies. Further, we generated and compared models using spherical and cartesian coordinate systems and monocular (Left or Right) or binocular data. All calibration routines performed similarly, with the best performance (i.e., sub-centimetre errors) coming from the naturalistic task trials when the participant is looking at an object in front of them. We found that spherical coordinate systems generate the most accurate gaze vectors with no differences in accuracy when using monocular or binocular data. Overall, we recommend one-minute calibration routines using binocular pupil data combined with a spherical world coordinate system to produce the highest quality gaze vectors.

## 1 Introduction

The majority of laboratory examinations of eye gaze are highly constrained and reliant on the assumption that gaze behaviors are task-invariant [11]. That is, many laboratory tasks do not reflect naturalistic behaviours. Common sense says that where someone is looking is dependent upon both eye and head movements [14, 28], meaning head position must be accounted for when calculating and analyzing gaze. Most studies investigating hand-eye coordination circumvent this problem by restricting head movements through the use of a chin rest [16]. In the real world, we are free to gaze at objects throughout our full field of view, or even anywhere in our 3D space, provided we can turn and move. But, in the lab, the areas the participant can interact with are typically severely limited, such as restricting gaze to a computer monitor or tabletop [27, 19]. Controlling for such environmental variables lets researchers ask specific questions about the motor and neural mechanisms that govern hand-eye coordination but fail to ask how gaze performs in natural settings. When collecting data outside of the laboratory, it is simply not feasible nor ecologically valid to restrict movement of the head or restrict gaze to the interaction with a limited amount of space. Additionally, real-world data tends to be much more difficult to process and analyze because of the permissive setting in which it is collected; free movement of the body is encouraged, as it more closely reflects natural behaviour.

Collecting data outside of the laboratory—or “in the wild”—is challenging [16]; determining fixations from dynamic bodies moving through dynamic worlds is a nontrivial problem to solve. A few studies have collected data while performing simple every-day activities [15, 13, 6, 29, 17]. For example, Land and Hayhoe [15] found that eye behaviours were similar across different use cases, such as during making a cup of tea or preparing a sandwich. They found eye movements could be broken down into four systematic categories: locating (the target), directing (the hands to the target), guiding (the hands during movement), and checking (if the condition has been satisfied). These general rules of interaction help inform us of potential systematic analyses that can be performed on the data. Data recorded “in the wild” also tend to be harder to parse into fractional chunks for analysis; Lappi [16] describes some of these common issues when collecting real-world natural gaze behaviours. In his review, Lappi suggests that complex eye movement behaviours are built from combinations of primitive eye behaviours such as fixations, saccades, and pursuits. These primitive building blocks can be used as indices to break complex tasks into digestible blocks that can be analyzed more similarly to controlled lab-based experiments.

Over the last decade eye tracking technology has become cheaper and easier to use. Traditional eye tracking headsets tended to be bulkier and required the head position to be fixed, whereas newer eye trackers such as the Pupil Labs Core [10] are more portable and do not require a fixed head. One common consideration of designing an eye tracking study is the time-consuming manual labour required for cleaning and analysis [17, 29, 32, 21]. Much of this manual labour is centred around two primarily video based categorization steps: 1) the cleaning of the pupil data, most of which is difficult to automate because of the nature of data quality from individual participants and 2) the assignment of fixations to objects in the world on the “world camera”, an outward facing camera attached to a head mounted eye tracker. This portion of analysis is so time consuming that many researchers will only analyze a subset of data rather than the whole [33, 30, 21]. For example, Parr et al. [21] were only able to analyze every third trial of their prosthetic hand-eye coordination task. A major concern is that a subset of data does not always represent the population-level statistics of the entire dataset—effects could be driven by outliers. Secondly, this leaves open the possibility of incorrect coding, leading to lower quality data that may contain additional errors, influencing statistical tests. Optimizing the volume of data analysis possible would have great benefits for statistical power and data reliability.

Motion capture (mo-cap) is a technique used to record human movement in 3D space [12] and, when combined and synchronized with eye tracking, offers a solution for automating real-world gaze analysis. Human movement science greatly benefits from this technology, as it allows for the quantification of movements during reach-to-grasp [3, 34] or reach-to-point [24, 23, 31] behaviours. Additionally, mo-cap technology comes in many forms, including infrared-based or the burgeoning field of markerless-motion-capture [18], both of which are typically capable of integrating with eye tracking headsets. Tracking gaze during movement grants insight into the strategies that different populations may use when completing the same task. For example, research into gaze strategies during reach-to-grasp behaviours has uncovered key strategic differences in normative [21] versus prosthetic arm users [32, 17, 8], where prosthesis users tend to move much slower, fixate longer, and do not “look ahead” to the intended target location after grasping.

While the synchronized collection of eye tracking and mo-cap data is not trivial, tools such as Lab Streaming Layer (LSL; SCCN [25]) have made this process much easier. However, once you have two synchronized data streams, it is not easy to determine where someone is looking based on raw data. Here we explore a technique requiring the experimenter to collect a separate eye-calibration data file, specifically for the purposes of building a model that will map head and pupil positions to a three dimensional (3D) gaze vector in a common world coordinate system. A 3D gaze vector is a line that extends from the head out into 3D space to predict where the participant is looking in world-space [26, 2, 20]. During these eye-calibration trials, participants are asked to focus on a tracked mo-cap marker (in our task on the tip of a calibration “wand”) as it moves through space, typically for about a minute. In the 2D eye tracking space, there does exist some guidelines and recommendations to generate gaze points. However, to our knowledge, despite the increasing number of studies that use 3D gaze vectors to assess behaviour, no standardized and very few recommended calibration routines exists. That is, how should you best move the tracked “wand”-marker through space? In addition, what data should be used to build predictive models, including which coordinate frame(s) or binocular / monocular data, depending on how pupil data were recorded.

With the goal of providing researchers interested in naturalistic tasks recommendations and guidelines for expected accuracy, we generated and assessed 3D gaze vector models from all possible combinations of: four different calibration routines, two coordinate frames, and three sets of pupil data inputs. Our results describe an approach that is capable of generating accurate (sub-centimetre and below one visual degree in the best case) 3D gaze vectors (GVs) using the position of the pupils and the 3D location of the participant’s head in space. To create the GVs, we use a linear regression algorithm to train models based on input pupil positions time-synchronized to the 3D location of a calibration wand. Then, we assess their spatial accuracy across a variety of data sets.

## 2 Methods

### 2.1 Equipment

Eye tracking data were collected using a Pupil Labs Core (200Hz; [10]) USB eye tracking headset. Lab Streaming Layer (LSL; [25]) was used to synchronize eye tracking and mo-cap data. The official Pupil Labs LSL plugin was used in conjunction with the Pupil Capture software to directly send data into the LSL datastream. Mo-cap data were collected using an OptiTrack mo-cap system (two systems were used throughout the study as the lab was upgraded: initially a 12-camera Flex 13 system, 120Hz; then a 14-camera Prime 13-W system, 200Hz). The OptiTrack systems were calibrated using the included Motive program to have a spatial accuracy of 0.1mm or less. A custom program was written in C# to pass frame data from the OptiTrack Motive application to the LSL datastream for synchronization. Rigid clusters of reflective markers were fixed to the participant and objects in the environment to track the position and orientation of the Head, Right Hand (centred approximately dorsally), Task Cart, Side Cart, Pasta Box (in Task data), and a Calibration Wand (in calibration data). Marker clusters were also fixed to the participant’s pelvis, trunk, upper arms, forearms, and left hand in as described by Boser et al. [1], but these data were not used in the current study. It is worth noting that theoretically any combination of eye tracker and mo-cap system could be used, provided they collect time series data as synchronized 2D pupil positions (in eye camera coordinates) and 3D marker position (in mo-cap).

### 2.2 Participants

Twenty-one undergraduate and graduate students from the Department of Psychology research pool at the University of Alberta participated in this study. All participants were right-handed, had normal or corrected-to-normal vision, and were naive to the tasks. Eight participants were collected using the OptiTrack Flex 13 system at 120 Hz, and 13 were collected on the OptiTrack Prime 13-W system at 200 Hz. One participant was removed due to recording errors (poor tracking quality), for a total of twenty participants. This study was approved by the University of Alberta Health Research Ethics Board under protocol Pro00087329 and ethical protocols were in adherence to the 1964 Declaration of Helsinki.

### 2.3 Procedure

Each test of data quality consisted of 3 sets of Calibration/Validation trials and 2 sets of 10 Task trials, proceeding in the following order:

1. Calibration/Validation set
2. Task set
3. Calibration/Validation set
4. Task set
5. Calibration/Validation set

Each Calibration/Validation set included four Calibration trials (one of each type described below) and one Validation trial presented in a pseudo-random order. Each Task set included 10 repetitions of the previously published Pasta Box task (see [29] for a full description of the task parameters). In short, the Pasta Box task requires the participant to move a rectangular box of pasta between three key locations: the Side Cart, the Green shelf, and the Blue shelf. In between each of the reaches, the participant must touch the Home position (see Fig. 3 and section 2.3.3 for a visual representation of the task and relevant spaces). Each trial takes approximately 15 seconds to complete. In total, participants performed 12 Calibration trials (3 repetitions of each of 4 types), 3 Validation trials and 20 Task trials. Not all participants had usable data for every trials; we discuss dealing with missing data and removal in section 2.4.1.

**Fig. 1.**
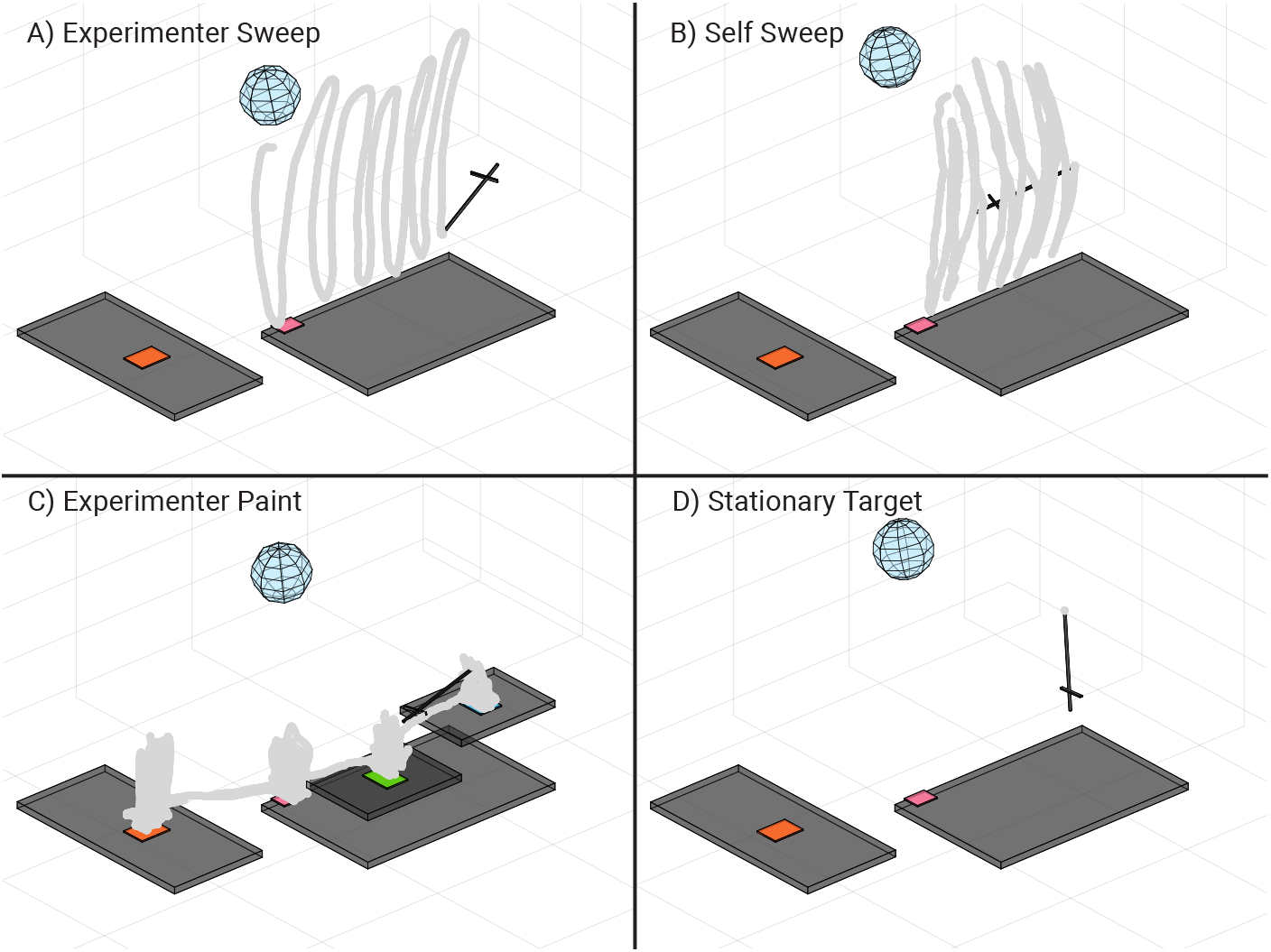
The Calibration routines used in the present study, with traces in grey showing example wand movements over time. Each routine takes approximately one minute to complete. The black inverted ‘t’ shaped object is the calibration wand used in all routines. In all quadrants, the blue sphere represents the participant’s head position, with the Orange target to the participant’s right and the wand being directly in front of the participant. A) The Experimenter Sweep (ES) routine. The experimenter stands to the participant’s left and waves the wand in s-shaped patterns through space, covering all three dimensions roughly equally (only up/down movements shown in figure). B) The Self Sweep (SS) routine. Identical in procedure to the ES routine, but the participant themselves carry out the wand movements. C) The Experimenter Paint (EP) routine. The experimenter stands to the right of the participant and moves the wand for approximately 15 seconds in small volumes at four locations relevant to the later Task trials: the Side Cart, the Home position, the Green Shelf and the Blue Shelf. D) The Stationary Target (ST) routine. The participant locks their gaze on the wand, which is fixed to the table. The participant moves their head up, then down, then centers, then left, then right (i.e., in the form of a cross), then rotates their head in swirl-like motions while maintaining fixation on the tip of the wand.

**Fig. 2.**
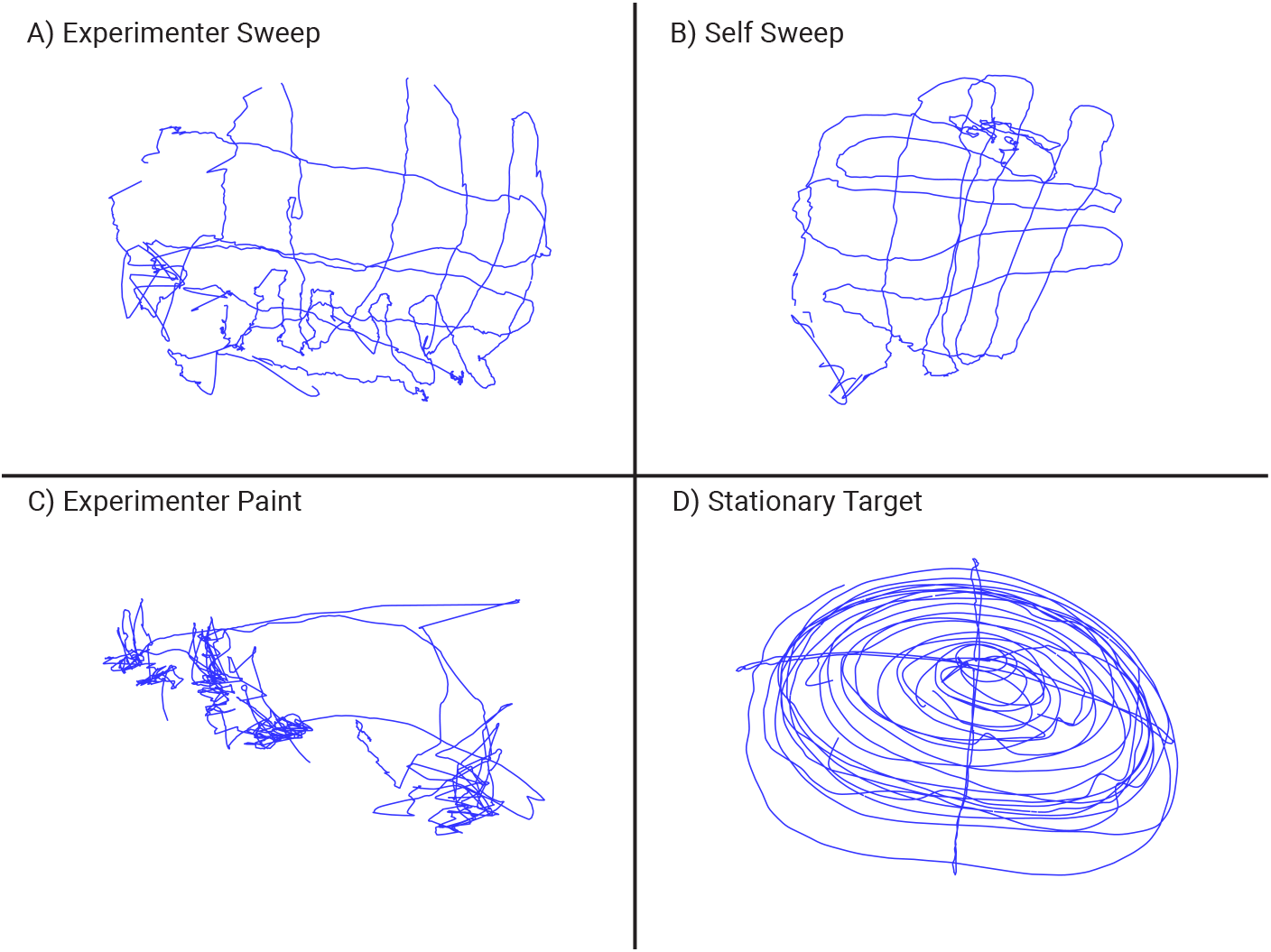
Example corresponding gaze patterns associated with each of the calibration routines. Pupil position from one eye is shown over the course of the entire calibration. A) The Experimenter Sweep (ES) routine: the gaze seems to be slightly jittery because the participant has to constantly adjust to the experimenter’s wand position. B) The Self Sweep (SS) routine: the gaze pattern is much more smooth, because the participant is moving the wand while simultaneously fixating on the tip. C) The Experimenter Paint (EP) routine: gaze locks to four different locations, which slightly overlap because the participant was free to move their head and likely tends toward central fixation on each location. D) The Stationary Target (ST) routine: the head is moved in a cross-like movement (up, down, centre, left, right) then in swirl-like movements for approximately one minute.

**Fig. 3.**
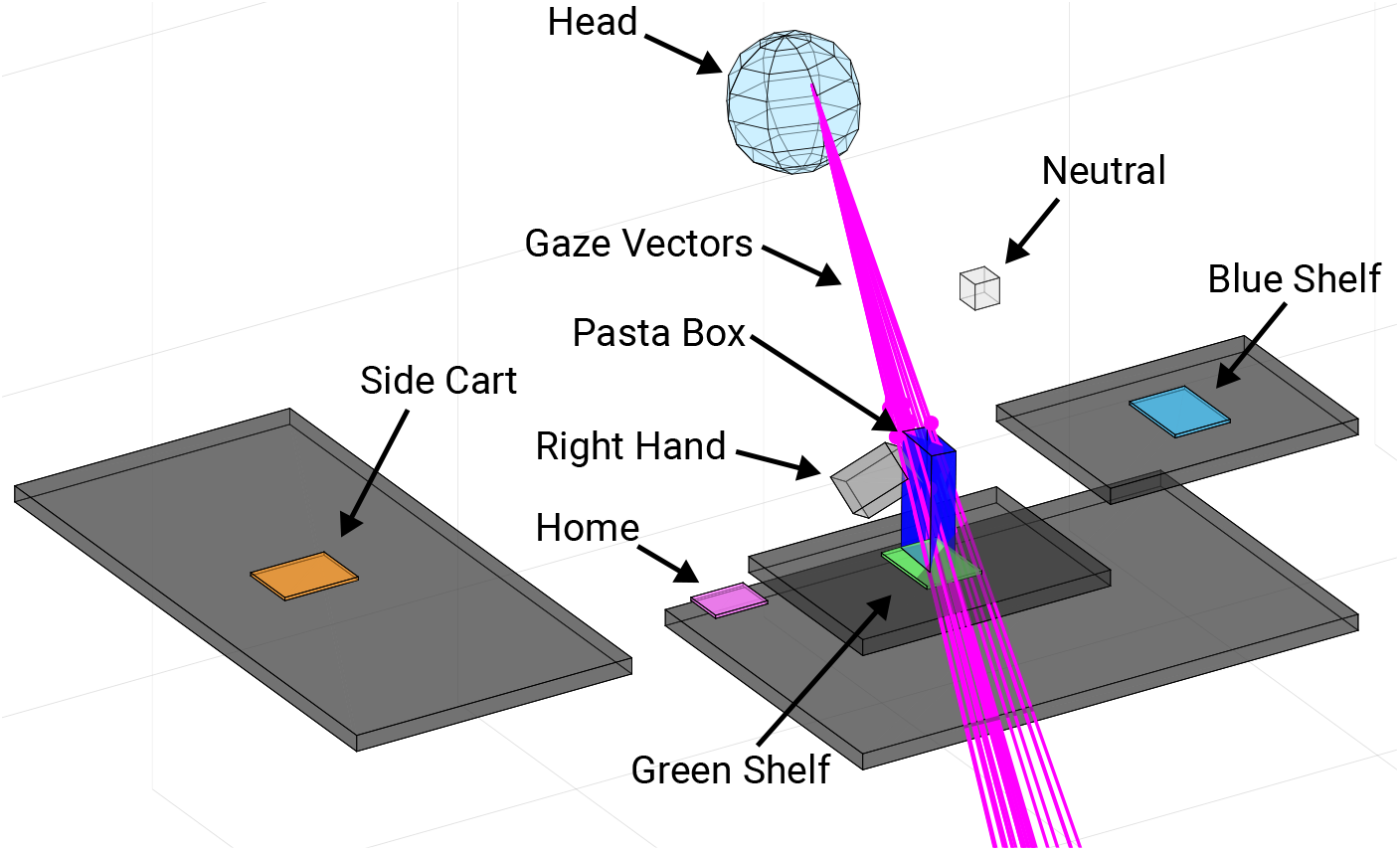
The locations, objects, and markers critical for all three tasks. The five locations are shown: Neutral, Side Cart, Home, Green Shelf, and Blue Shelf. For the Pasta Box task, the participant moved the box from location to location (see 2.3.3). The Head rigid body was used to determine the origin of the resulting gaze vectors. The Right Hand’s velocity profile was used to determine when the participant picked up or dropped off the pasta box (see 2.4.3). All 72 gaze vectors generated are shown in pink, with most being close to the target object (pasta box), and some performing rather poorly.

#### 2.3.1 Calibration Trials

Participants were asked to track the position of a single spherical mo-cap marker (14 mm diameter) with their eyes for about one minute per trial. The participant could move their head freely while tracking the marker. The marker was placed at the tip of a 40 cm wand which moved through the task space in one of four Calibration routines:

1. *Experimenter Sweep (ES):* The experimenter moved the wand in slow S-shaped curves along each of the room-coordinate axes (parallel to floor, left/right, parallel to floor in/out, parallel to wall up/down).
2. *Self Sweep (SS):* Replicating ES but with the participant holding the wand and replicating the movements.
3. *Experimenter Paint (EP):* The experimenter moved the wand to each of the relevant locations in the Pasta Box task (minus Neutral, see below) and explored small (10-20 cm in each dimension) volumes at these locations.
4. *Stationary Target (ST):* The wand was fixed to the table directly in front of the participant (~60 cm away), who was asked to maintain fixation on the wand-tip

while nodding their head up and down, returning to centre, then turning it left and right, then rotating it in a clockwise then counterclockwise spiral.

The intention for each of these trials was to create calibration routines with a diversity of different coverages in terms of both task and pupil-position space (see Fig. 1 for the wand movements, and Fig. 2 for example corresponding pupil positions).

#### 2.3.2 Validation Trials

Participants were asked to fixate on 5 stationary targets (see Fig. 3 for locations) presented at Task-relevant locations for ~5 s, in a specific sequence, and at least 2 times each. An auditory beep signalled the start of the first fixation and beeped every 5 seconds thereafter to signal a switch to the next Task-relevant location in this order:

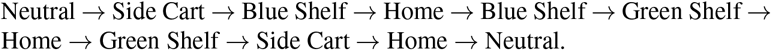

This order of 11 fixations mirrors the order these locations are visited during the actual Task trials.

#### 2.3.3 Task Trials

The set-up for the Pasta Box task is shown in Fig. 3^1^. Participants began each Task trial with their hand on the Home position and their eyes fixating on the Neutral target, marked by a mo-cap marker. A beep then cued them to initiate an object interaction sequence consisting of three movements:

1. Reach and grasp the Pasta Box at the Side Cart, move it to Green Shelf then return hand to Home;
2. Reach and grasp the Pasta Box at Green Shelf, move it to Blue Shelf then return the hand to Home;
3. Reach and grasp the Pasta Box at Blue Shelf, move it to the Side Cart then return the hand to Home. At the end of the task the participant also returns their gaze to the neutral marker.

The task was demonstrated to each participant visually. The participant was given as many practice trials as they felt necessary to be comfortable with the sequence of movements.

### 2.4 Data Processing

#### 2.4.1 Pre-processing

Mo-cap data were exported from Motive and run through custom MATLAB scripts to check for marker name consistency and remove residual sections of noisy data (marker displacements of more than 5 mm between frames, and islands of data less than 100 ms in duration). Mo-cap and eye tracking data were then synchronized to the mo-cap frame rate using the common timestamps in the LSL datastream files. The combined data were imported into our custom software platform for integrated analysis of eye and mo-cap data; the Gaze and Movement Assessment Tool (GaMA; [32]). Within GaMA, raw pupil position data was cleaned by: 1) Removing any data points outside of pupil camera bounds (<0 or >1); 2) Removing any data points more than 4 standard deviations away from the mean position; 3) Removing any data points with velocities greater than 6 (meaning the pupil was travelling across the entire camera 6 or more times per second). After this removal, any gaps < 50 ms were filled using the *inpaint_nans* [5] function in MATLAB then, any remaining islands of data < 50 ms were deleted. Finally, the pupil data were filtered in MATLAB using a 4^th^ order zero-lag low-pass Butterworth filter with a cutoff frequency of 10 Hz. A 10 Hz cutoff was chosen because the demands of the tasks do not depend on eye dynamics with movements more than 10 times / second. Also within GaMA, the mo-cap data were filtered using a 4^th^ order zero-lag low-pass Butterworth filter with a cutoff frequency of 6 Hz. Rigid bodies, represented as both a position and rotation, were defined using the clusters of markers attached to the participant’s head and hand, as well as objects in the environment. For the Task trial data, virtual objects were also created to represent the position, orientation and extent of the objects in the environment (Task Cart, Side Cart, Pasta Box).

#### 2.4.2 Gaze Vector Modelling

The cleaned eye and motion data were then used to generate predictions of the direction the participant was looking in 3D space, or “gaze-in-world” vectors, herein referred to as GVs. The process of generating a single GV consists of two steps:

1. Generate eye gaze models using data from a specific Calibration trial
2. Use the eye gaze models to predict the GV direction at each frame in a given trial

In step 1, Calibration data are used to fit three eye gaze models. Each model takes pupil position data as input and predicts a single coordinate of the 3D gaze fixation point relative to the Head rigid body coordinate system in the 3D mo-cap space. For example: one model might use pupil position data to predict only the x-coordinate of the fixation point relative to the head, a second, separate model would be used to predict only the y-coordinate, etc. Each eye gaze model was generated using the built-in MATLAB function `fitlm` with the ‘`quadratic`’ model specification and robust fitting using the ‘`bisquare`’ weight function. i.e.:

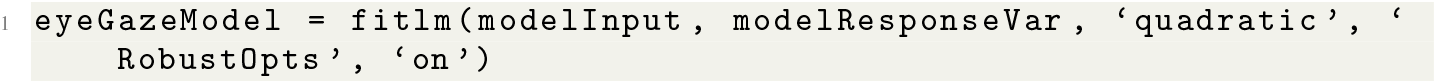

In this study we explored three options for model input (pupil input data from right eye only [x_r_,y_r_], left eye only [x_l_,y_l_], or binocular data [x_l_,y_l_,x_r_,y_r_]), as well as two options for expressing the fixation point relative to the Head coordinate system (Cartesian [x,y,z] coordinates, or Spherical [r, *θ*, *φ*] coordinates). We anticipated that using the Spherical coordinate system would increase accuracy of the GV direction because it isolates depth of fixation to the ‘r’ model, whereas in Cartesian, all three models are influenced by depth of fixation.

In step 2, once the eye gaze models were generated for a given Calibration trial and set of parameters (left/right/both eyes × cartesian / spherical coordinate system), they were used to predict the coordinates of the fixation point relative to the head at each frame in a given Calibration, Validation, or Task trial. The known transformation between the Head rigid body coordinate system and global mo-cap coordinate system is then used to calculate the position of the fixation point relative to the global coordinate system. The GV is represented by the line originating at the head rigid body origin (mid forehead), passing through the fixation point, extending infinitely forward and away from the head in the direction of the fixation point (see Fig 4). It is important to note that only the direction of the GV was used in subsequent analysis, the distance from the head to the predicted fixation point was not considered.

**Fig. 4.**
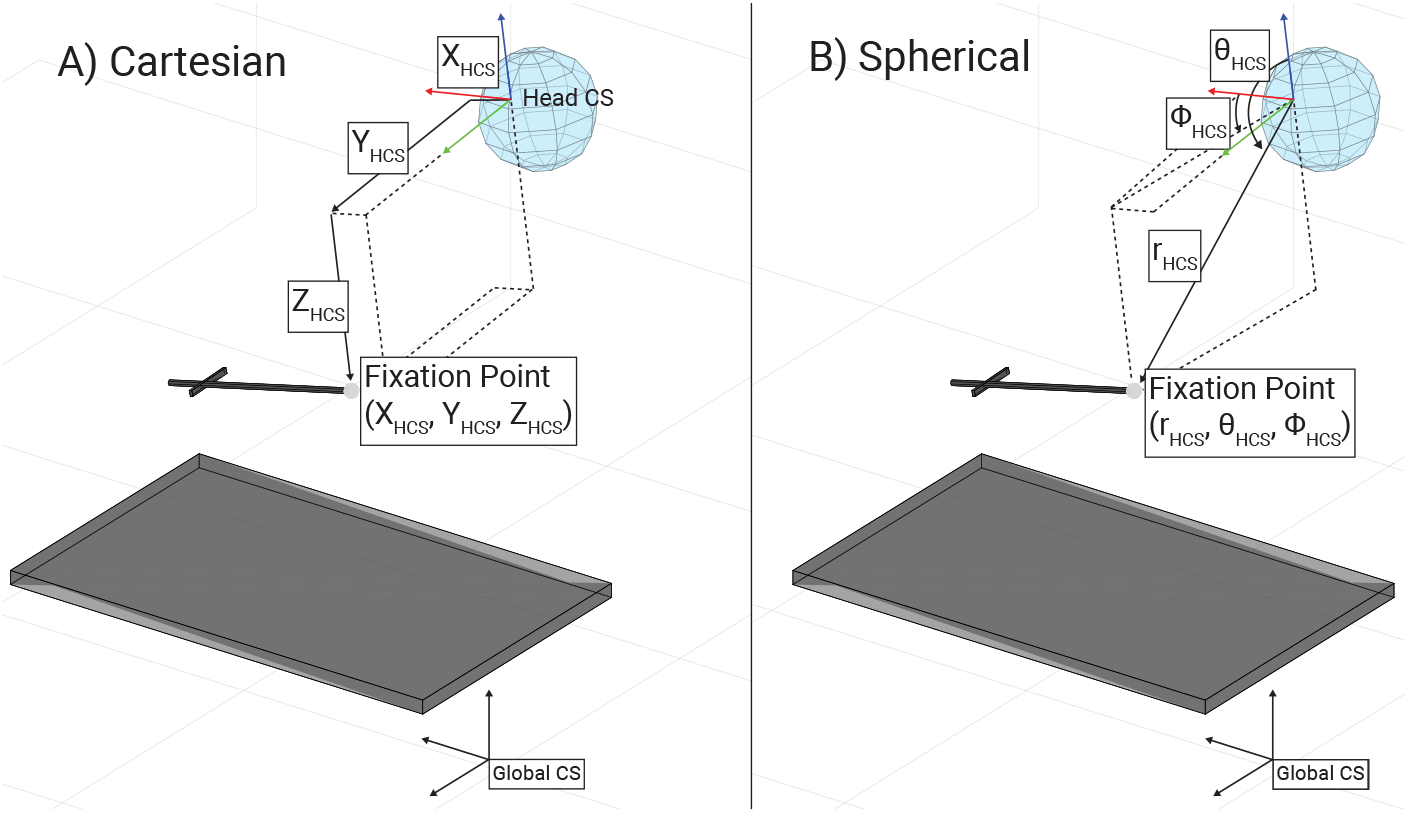
A visual demonstration of the differences between the Cartesian and Spherical coordinate systems used. The tip of the black wand is the gaze fixation point in both coordinate systems. A) The Cartesian coordinate system: coordinates are represented by coordinate triplets of [x,y,z]. Here, the wand tip is only represented by its offset from the origin. A consequence is that the depth of the wand is embedded in all of the dimensions. B) The Spherical coordinate system: coordinates are encoded as triplets of [r, *θ*, *φ*]. Here, the wand tip is described in terms of the angular and euclidean distance from the head. As a consequence, euclidean distance (i.e., depth) can be sequestered to a single coordinate. As our goals did not require depth estimates, we were able to train our models without the depth information.

#### 2.4.3 Dependent Measures

We conducted separate analyses to assess GV accuracy in each of the three types of data collected (Calibration, Validation and Task). For each analysis, we collapsed across the 3 repetitions of a given Calibration type by finding which of the repetitions performed the “best” on that data. This involved eliminating abnormally poor GVs (those whose average distance from the target of analysis were 30 cm or more away), then taking the remaining GV with the lowest average distance to targets (see below). One participant was removed from the Calibration dataset, three from the Validation dataset, and two from the Task dataset because of average errors above 30cm. An advantage of this approach was this it allowed participants to be included for analysis that may have had errors in recording one repetition of Calibration data. As linear distance error does not account for the perspective of the participant, we also calculated the visual angle error simultaneously for each trial. The visual angle error accounts for the distance between the subject’s eyes and the target object.

##### Calibration Trials

The dependent measure for Calibration trials was the mean 3D distance between a given GV and the Wand Tip over the entire trial. To reiterate from above, distances were always calculated as the minimum distance between the GV line and the Wand Tip point, meaning the depth of fixation along the GV was not a determinant of accuracy.

For each of the 12 Calibration trials we generated all 72 possible GVs (12 Calibration files × 2 Coordinate systems × 3 pupil data inputs). For each coordinate system and input data combination, we compared the three repetitions of GVs of the same type (e.g., across the 3 ES GVs that were Cartesian and with Both eyes) across all 12 Calibration trials to find the one that performed the best. Note that trials were not tested on the data used to train the model. First, we eliminated GV outliers (those with GV to Wand Tip distances > 30 cm), then we took the median performance for each of the 3 repetitions across the remaining Calibration trials. The repetition with the lowest median performance was then selected as the Best and used for the remaining analyses.

##### Validation Trials

The dependent measure for Validation trials was the mean 3D distance between a GV and each of the Task Relevant target locations. To extract these Task Relevant target looks, we isolated 1 second epochs of stable-gaze data in the 5 s between the cueing-beeps. For example, between the first and second beep, participants were instructed to look at the Neutral target. Within this 5 s window, we use a modified moving mean algorithm to find the 1 s of data where 1) there was at least 50% of detected pupil and 2) the L or R pupil data has the lowest velocity. This process generates 11 stable-gaze epochs, three for the Home location and two each for the remaining four Task-relevant locations. For each of the five Task-relevant locations, the reported distance is the mean over these stable-gaze epochs.

Similar to the Calibration trials, but across the 3 Validation trials, we selected the best GV within a set of repetitions by comparing their performance across the 5 locations. First, we eliminated outliers (over 30 cm mean distance from any location), then took the one with the minimum median distance across all Validation trials and the five locations as the best GV.

##### Task Trials

The dependent measure for the Task trials was the mean 3D distance between a given GV and the nearest bounding box face of the Neutral (4 cm cube) or Pasta Box (9 x 4 x 18 cm; see Fig. 3) object at specific locations and times during the interaction task. Eye gaze behaviour is well understood for this task, as described in Lavoie et al. [17] and Williams et al. [32]. Following the same procedure as in this earlier work, each task trial was segmented into specific movements and movement-phases (Reach, Grasp, Transport and Release) using detailed procedures described elsewhere [17]. For this analysis, we isolated looks toward the Neutral marker at the start of the trial and looks toward the Pasta Box each time it was being grasped (just prior to object pickup) and released (just after object dropoff). Previous work using this identical task shows that there are fixations to these objects around these times on almost every trial [17]. These were single frame events that occurred once (for the look to Neutral) or twice (for the looks toward the Pasta Box at the Side Cart, Green Shelf and Blue Shelf locations) per location. Distances to locations with two looks were averaged.

Similar to the Calibration and Validation trials, across the 20 possible Task trials we selected the best GV within a set of repetitions by comparing their performance across the 4 locations (note no interactions occurred at the Home location so it was not included in the Task trial analysis). First, we eliminated outliers (over 30 cm mean distance from any object), then took as the best GV the one with the minimum median distance across all Task trials and the four locations.

## 3 Results

Statistical analysis was performed in JASP 0.16.1 [9]. Repeated-measures ANOVAs (rmANOVAs) were used to analyze the three trial types, which used the same participant pool but were statistically independent from one another. We conducted statistical analysis on two independent sets of data: a linear distance error (centimetres) and a visual angle error (degrees) to account for distance. All results below are reported with the linear distance error (LD) first and visual angle error (VA) second. We opted to use a conservative statistical approach, correcting α for the number of tests run in each family as described by Cramer et al. [4]. Each of the trial types were considered a family for this analysis. All *p* values were Greenhouse-Geisser corrected if sphericity was violated and more than two levels existed in the factor.

### 3.1 Factors for rmANOVA

For clarity, here we lay out all of the factors and their levels input into each rmANOVA. The Coordinate factor describes the type of coordinate frame used. The Eye factor describes whether monocular (left or right) or binocular data were used. The Calibration factor describes the routine when testing on Calibration data. The PredictedCalibration factor describes what data were *input* into the model to calculate the errors. The Location factor describes the specific location that the participant was to interact with.

#### Levels

All Coordinates had two levels: Cartesian and Spherical. Eye had three levels: Right, Left, and Both. Calibration had four levels: ExperimenterSweep, Paint, Self, and Stationary (see 2.3.1). PredictedCalibration had four levels: ExperimenterSweep, Paint, Self, and Stationary. Location had five levels in the Validation trials: Neutral, SideCart, Home, GreenShelf, and BlueShelf but only four levels in the Task trials: Neutral, SideCart, GreenShelf, and BlueShelf (see Fig. 3).

### 3.2 Calibration Trials

Here, we ran an rmANOVA on a 2 (Coordinate) × 3 (Eye) × 4 (Calibration) × 4 (PredictedCalibration) design.

A significant main effect of Coordinate was detected (LD: F(1,1) = 72.984, *p* < 0.001, *η*^2^ = 0.024; VA: F(1,1 = 100.586, *p* < 0.001, *η*^2^ = 0.020), where a model generated using Spherical data had lower error than with Cartesian data (see Fig. 5). A significant main effect of Calibration was detected (LD: F(1,1.971) = 11.894, *p* < 0.001, η^2^ = 0.050; VA: F(1,2.058) = 12.393, *p* < 0.001, η^2^ = 0.047), with Stationary data on average performing best. A significant main effect of PredictedData was detected (LD: F(1,1.870) = 8.660, *p* < 0.001, η^2^ = 0.128; VA: F(1,2.294) = 6.995, *p* < 0.001, η^2^ = 0.119), with Stationary data being predicted more accurately in a Spherical coordinate system. A significant Coordinate × PredictedData interaction was detected (LD: F(1,1.259) = 18.377, *p* < 0.001, η^2^ = 0.015; VA: F(1,1.215) = 24.979, *p* < 0.001, η^2^ = 0.016), where Stationary data were the hardest to predict when predicted by non-Stationary models, but performed well when predicted by a Stationary Calibration model. A significant Coordinate × Calibration interaction was detected (LD: F(1,2.589) = 20.498, *p* < 0.001, η^2^ = 0.012; VA: F(1,2.114) = 26.167, *p* < 0.001, η^2^ = 0.010), with Stationary data again being hard to predict, unless it is predicted by a Stationary model. A significant Coordinate × Calibration × PredictedData interaction was detected (LD: F(1,4.338) = 4.942, *p* < 0.001, η^2^ = 0.006; VA: F(1,3.834) = 7.644, *p* < 0.001, η^2^ = 0.006), driven by the performance of Stationary data on non-Stationary Calibration models (see Fig. 5B and D).

**Fig. 5.**
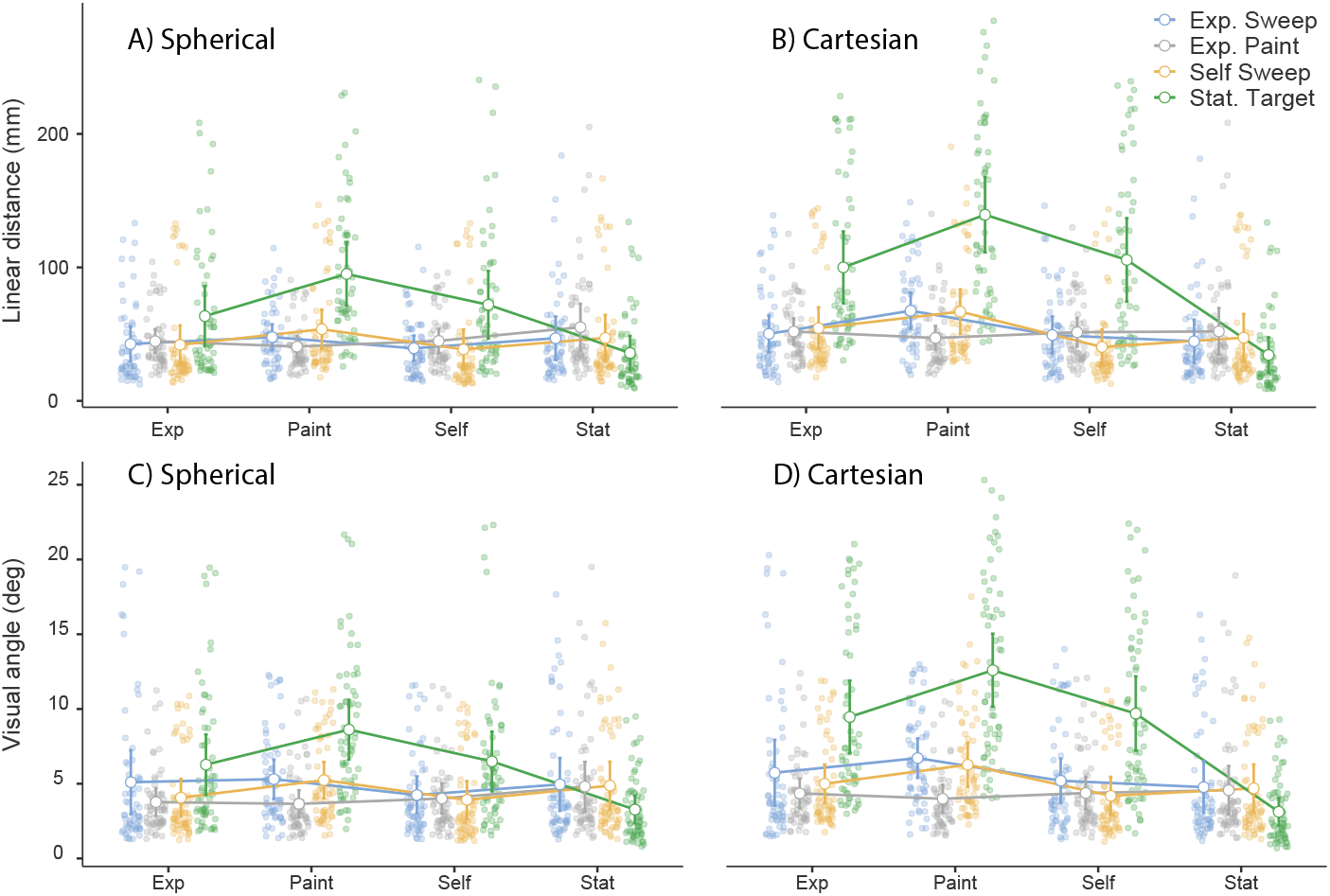
Plots showing the average performance of the models in linear distance (LD) and visual angle (VA) errors for the Calibration trials. The legend in the top right denotes which Calibration data was used during assessment. The top row are plots of LD errors (in mm) and the bottom row shows VA error (in degrees). The left plots are Spherical coordinate data, and the right plots are Cartesian coordinate data. 95% confidence intervals are used around each mean, with the observed scores that make up that mean plotted as translucent points. In all plots, the X axis denotes the Calibration routine used to train the model. A) Mean error of the Spherical LD models tested. Error is shown in mm at each location. B) Mean error of the Cartesian LD models tested. Error is shown in mm at each location. C) Mean error of the Spherical VA models tested. Error is shown in degrees at each location. D) Mean error of the Cartesian VA models tested. Error is shown in degrees at each location.

All other tests were either not significant or were rejected because they did not meet Cramer’s adjusted α criterion [4].

### 3.3 Validation Trials

We used an rmANOVA on a 2 (Coordinate) × 3 (Eye) × 4 (Calibration) × 5 (Location) design.

A significant main effect of Coordinate was detected (LD: F(1,1) = 25.928, *p* < 0.001, η^2^ = 0.008; VA: F(1,1) = 33.716, *p* < 0.001, η^2^ = 0.003), where a model generated using Spherical data had lower error than with Cartesian data. A significant Coordinate × Calibration interaction was detected (LD: F(1,2.501) = 7.838, *p* < 0.001, *η*^2^ = 0.006; VA: F(1,3) = 7.274, *p* < 0.001, *η*^2^ = 0.002), where the Spherical models tended to outperform Cartesian models, except when testing on Stationary data (see 6B).

All other tests were either not significant or were rejected because they did not meet Cramer’s adjusted α criterion.

### 3.4 Task Trials

For the Task data, we were concerned with the performance of the GVs on real-world data. Here, we ran an rmANOVA on a 2 (Coordinate) × 3 (Eye) × 4 (Calibration) × 4 (Location) design.

A significant main effect of Coordinate system was detected (LD: F(1,17) = 21.475, *p* < 0.001, η^2^ = 0.008; VA: (F(1,17) = 18.748, *p* < 0.001, η^2^ = 0.006), where Spherical models had lower errors than Cartesian models (see Fig. 7). A sig-nificant main effect of Location was detected (LD: F(1,3) = 37.102, *p* < 0.001, η^2^ = 0.202; VA: F(1,3) = 20.550, *p* < 0.001, η^2^ = 0.083), where the SideCart location was the most difficult to predict, resulting in the highest errors overall (see Fig. 7A). A significant Coordinate × Calibration interaction was detected (LD: F(1,3) = 7.396, *p* = 0.001, η^2^ = 0.006; VA: F(1,3) = 11.177, *p* = 0.001, η^2^ = 0.004), with Spherical data outperforming Cartesian data in all cases except for Stationary data.

**Fig. 6.**
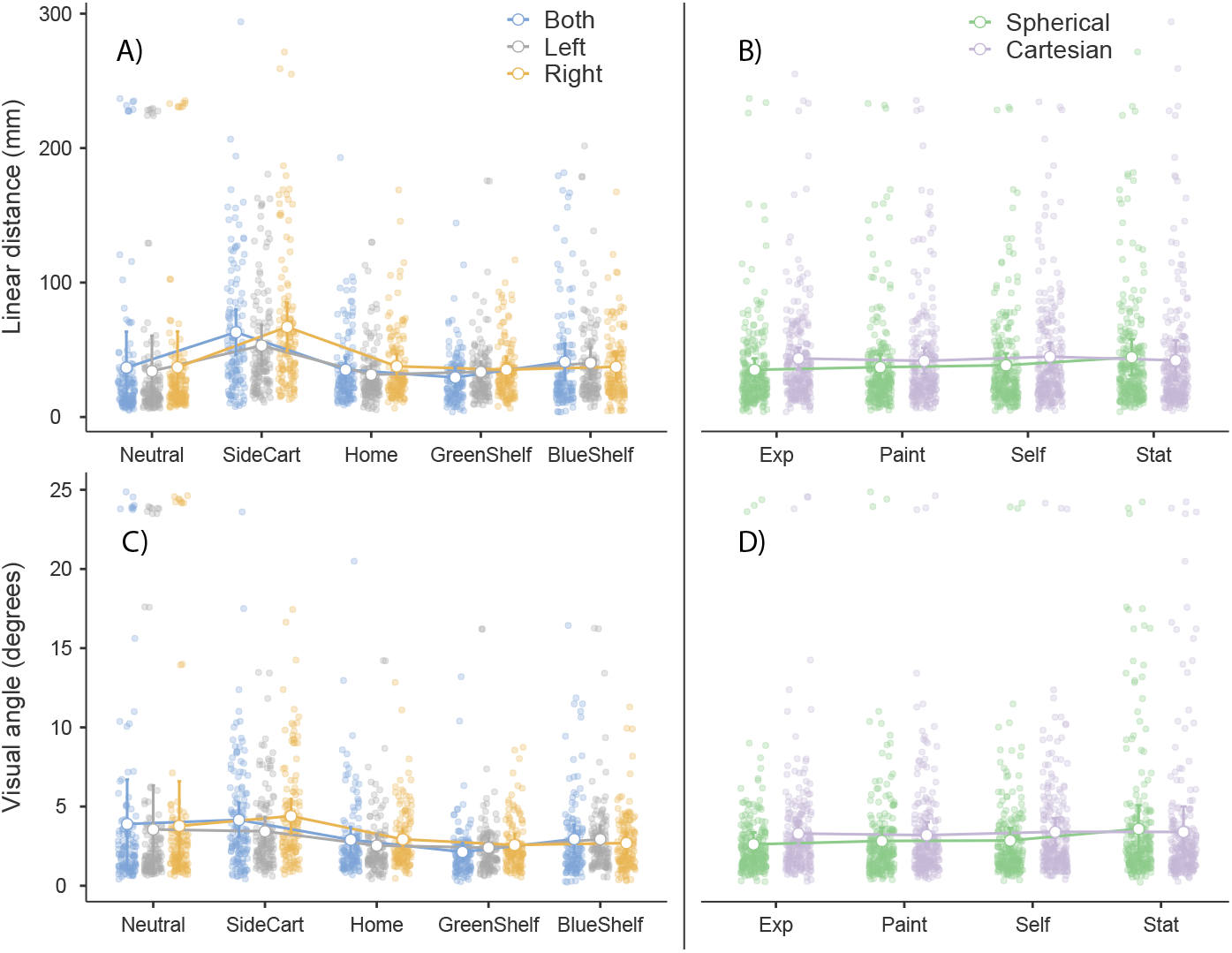
Plots showing the average performance of the models in linear distance (LD) and visual angle (VA) errors for the Validation trials. The plots on the left side are the average LD and VA errors (A, C) for each location in the Validation task (X axis). The legend here indicates what type of pupil input data was used. The plots on the right side the average LD and VA errors (B, D) for each of the Calibration types used as inputs to the models (X axis). The legend in the top right indicates whether a Spherical or Cartesian model was used. 95% confidence intervals are used around each mean, with the observed scores that make up that mean plotted as translucent points. A) Mean error generated at each of the locations (along the X axis) is shown for each type of Eye data used. Error is shown in millimetres at each location. B) Mean error generated each of the Calibration routines used is shown for Spherical and Cartesian models. Error shown in millimetres for each Calibration routine. C) Mean error generated at each of the locations (along the X axis) is shown for each type of Eye data used. Error is shown in degrees at each location. D) Mean error generated each of the Calibration routines used is shown for Spherical and Cartesian models. Error shown in degrees for each Calibration routine.

**Fig. 7.**
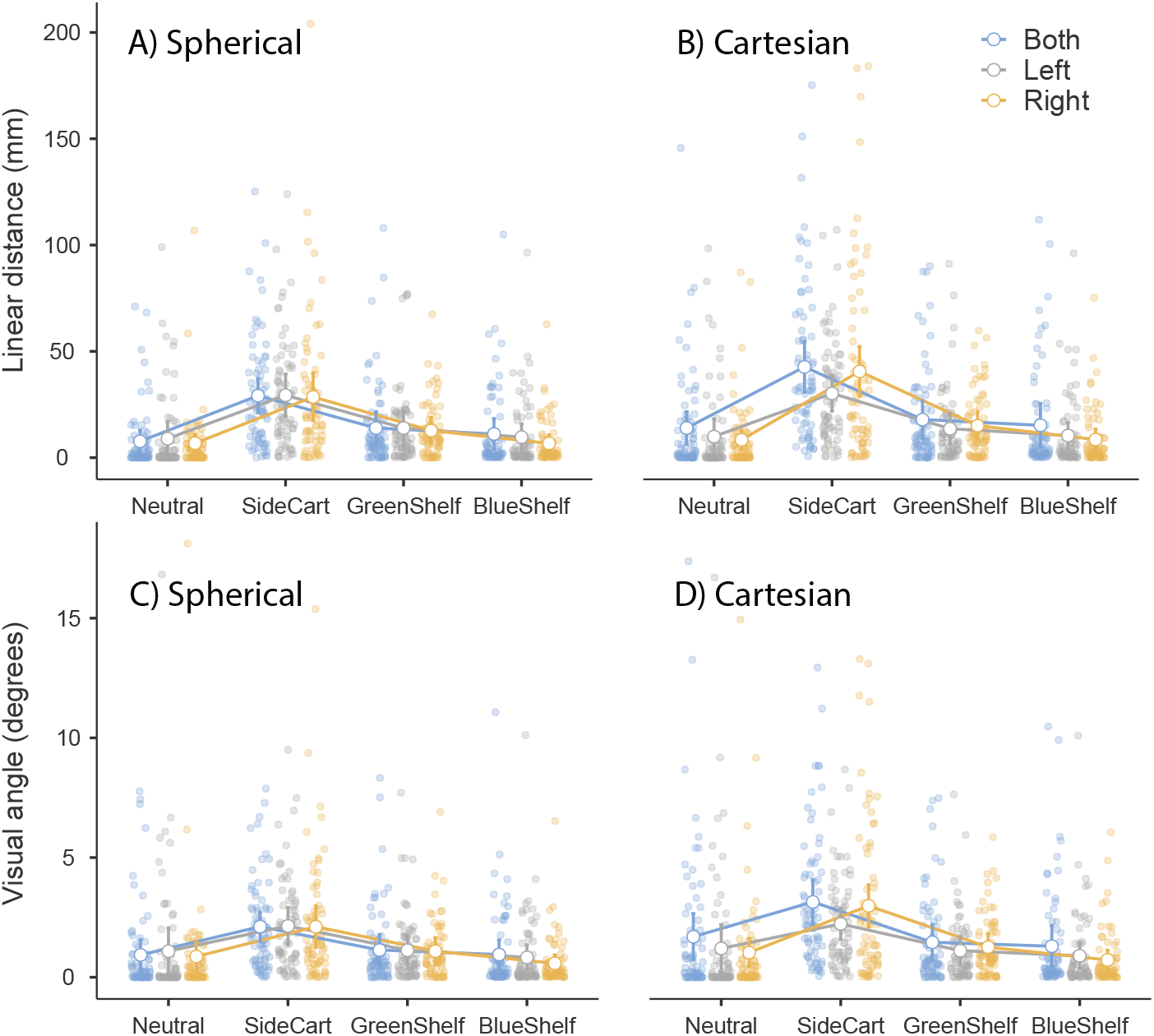
Plots showing the average performance at each location, split by the pupil input data in linear distance (LD) and visual angle (VA) errors for the Task trials. 95% confidence intervals are used around the mean points. The X axis denotes the Location during the Pasta Box task for all plots. A) Mean LD error for a Spherical coordinate system at each Location. Note that errors are below a centimetre when the participant is fixating on the Neutral marker (i.e., directly in front) or at the pasta box on the Blue Shelf. B) Mean LD error for a Cartesian coordinate system at each Location. C) Mean VA error for the Spherical coordinate system at each Location. Note that errors are below one degree of visual angle when fixating on either the Neutral marker or at the pasta box on the Blue Shelf. D) Mean VA error for a Cartesian coordinate system at each Location.

All other tests were either not significant or were rejected because they did not meet Cramer’s adjusted α criterion.

## 4 Discussion

Here we describe a method for generating 3D GVs using combined eye tracking and mo-cap data. We achieve this by collecting Calibration data where the eyes are continuously fixated on a tracked mo-cap marker, and using it to train a set of linear models to predict the 3D coordinates of the gaze fixation point. Within this method we explored four different Calibration routines (ES, SS, EP, ST), three options for eye input into the model (binocular, left, right), and two options for model coordinate system (spherical and cartesian).

We describe a set of four one-minute calibration procedures and their performance relative to one another in three different, but related analyses. All Calibration procedures were similar in that the goal of each was for the participant to fixate on a specific mo-cap marker for the duration of the procedure. We propose a simple model-based assessment (MATLAB’s `fitim` function) that allows us to give a recommendation for the best Calibration procedure based on the average GV error from a known location. First, we assessed a model trained on a Calibration routine’s eye and mo-cap data calculating its error when testing on all other Calibration procedures’ input data. Second, the participant completed several Validation trials. During these trials, the participant fixated on different areas of interest for long (~5 s) periods of time to effectively emulate eye gaze behaviour during our Pasta Box task [17, 1, 32, 29], allowing us to assess error at each area of interest used during the task. Finally, participants performed a real-world task where they were instructed to perform the described Pasta Box task. Here, we demonstrate that our analysis techniques extend to data that were not recorded for the express purpose of being put through this analysis pipeline. That is, we can assess real-world task data and calculate performance metrics to best determine which Calibration procedure to use.

With respect to the type of coordinate system used to predict the gaze fixation point relative to the head, the results demonstrate that using a Spherical coordinate system generally results in a GV with a more accurate direction than using a Cartesian coordinate system as well as lower error overall. This result is aligned with our prediction, as we expected that the depth of fixation would be difficult to model based on pupil position data alone. It is worth noting that the reduction in error appeared to be systematic across all Calibration routines used—Spherical outperforms Cartesian. Although we did not assess gaze depth in the current study, when a Cartesian coordinate system is used, both the depth of fixation and direction of gaze are partially represented in all three eye gaze models (the x, y, and z coordinates). Whereas, using spherical coordinates confines the depth to one model (the ‘r’ coordinate) which does not impact the direction of the GV. In future work we intend to further explore the accuracy of the depth of the fixation point. However, the present work indicates that when only the direction of gaze is of interest, a spherical coordinate system should be used to generate GVs. When comparing each Calibration procedure on their ability to predict Calibration data, all models perform relatively well, with Stationary performing the best. However this is driven by the fact that all Calibration procedures (excluding Stationary) appear to have a difficult time predicting Stationary data. The Stationary routine was intended to allow the eyes to explore the maximal range of trackable pupil-space (see Fig 2D) while the actual target remained constant in space, potentially leading to a more robust model. The Stationary Calibration takes advantage of the compensatory vestibular-ocular response (VOR; [35]), in that the eyes and head move, but the gaze target remains static. The approach for the Stationary Calibration is one of quantity over quality; during the Stationary routine, data is collected from almost all accessible areas of the pupil, but not a lot of time is spent at each location nor are many of these locations generally useful during the Pasta Box task.

While it may be tempting to conclude that Stationary performs best overall, the data actually collected during Validation and Task trials do not reflect this same level of pupil space exploration. It is also important to consider that data collected during actual trials do not typically result in the pupil being located in positions on the eye consistent with the Stationary routine. Therefore, despite the advantage that the Stationary routine appears to show for Calibration data, the fact that it did not perform better during the more ecologically valid Validation and Task trials leads us to recommend using a Calibration routine that reflects the dynamics of eye exploration necessary during task completion. Anecdotally, explaining the Paint Calibration procedure was the simplest to perform and is extensible to any task, while the Experimental Sweep procedure was the easiest to keep consistent between sessions. Therefore, one of these two would be our recommendation for ease and consistency without sacrificing performance for ecologically valid data.

The Validation task was designed to mimic the behaviours that occur during a typical Pasta Box trial while still giving control over where the participant is looking and when. During a real trial, it is much more difficult to intrinsically know where the participant should be looking. These results are in line with the Calibration results, suggesting that Spherical coordinates result in more accurate GVs. One of the challenges the model faces is when the participant turns to fixate on the Side Cart, which results in higher error. Side Cart error appears to be worse when using data from Both pupils, and performs best when using monocular data, notably from the Left eye. One possible explanation for this is that the Left eye is always in view of the cameras when fixating on the target at the Side Cart, whereas the Right eye is potentially lost for a short duration. It is possible that using a ‘hybrid’ approach with monocular pupil data input, constantly switching to the ‘better’ eye, could result in superior performance. However, this is to be investigated and cannot currently be stated for certain. Regardless, it does suggest that collecting data from both eyes gives the most flexibility and opportunity to maximize data quality across sessions and even within a task. The Validation dataset functions as a ‘sanity check’ to ensure that the performance of the model is at least in line with our expectations: instead of tracking a moving marker (e.g., the wand during Calibration trials), the participant is fixating on a single static marker at a Task-relevant location. Performance appears to be similar to the Calibration trials analysis, suggesting the Validation dataset has done its job.

The Task results demonstrate that performance of the model has been effective on real-world data using a well-documented task [29, 17, 1, 32]. Previous work has shown that normative participants tend to fixate on the object they are about to interact with (or about to stop interacting with) for several hundred milliseconds [6, 7, 22]. Assessing performance on a real-world task is challenging because the behaviours of the participant are not controlled beyond simple verbal instructions (e.g., pick up the pasta box and move it to a new location) or visual demonstrations. However, we can use the principals described by Lappi [16] and Hayhoe [6] to find points in time when we expect the participant to initiate a reaching behaviour, such as a fixation on the object to be interacted with. With the identified fixation, we assessed error at this time point as the minimum distance between the 3D GV and the Pasta Box. Overall, performance of the model looked good; errors were remarkably low (see Fig. 7A & C). The average error for a Spherical coordinate system was below a centimetre and under a degree of visual angle for Task trials. We were surprised to find that error was lowest in the Task trials as they were the least-controlled in terms of participant instruction. However, when the participant turned their head, the error was significantly higher than at other areas.

Currently, there do not seem to be any standardized calibration procedures that also allow for the assessment of performance during real-world task use. Here we show a methodology that allows anyone with access to an eye tracker that outputs pupil locations in 2D space and a motion tracker in 3D space to generate GVs that can have as low as sub-centimetre error. While we did not find that any particular Calibration routine’s data significantly outperformed any other, we found that using a Spherical coordinate system generated significantly less error on average when compared to a Cartesian coordinate system. Further, we suggest using a calibration routine that reflects the actual behaviours of the participant during task completion. For example, if the task involves looking at and reaching towards specific areas, a calibration routine that includes eye and hand movements towards those locations should generate higher quality models, or at minimum match the task demands and therefore be easier to employ.

## 5 Conclusion

We found that, when recording synchronized eye and mo-cap data for the purpose of producing accurate 3D gaze vectors, there are a few useful rules of thumb:

1. For fixations to real objects positioned in front of participants, gaze vectors generated using this approach will result in an average error of about 1-2 cm. If within peripersonal space (around 60 cm distance), this corresponds to about 1 visual degree.
2. A spherical coordinate system will on average produce more accurate gaze vectors (when depth is not considered).
3. Locations that require a head turn typically result in an accuracy falloff, adding about 2-3 cm of error in our data.
4. The best way to minimize error is to ensure quality data by making sure the eye tracker is properly fitted and the cameras are getting sufficient coverage of the eyes.
5. Binocular data, while not always the most accurate, gives the option to use either or both of the eyes when generating gaze vector models.
6. The calibration routine used should reflect the locations in space that the participant will be interacting with. More data is not always better.

## Data Availability and Open Practices

All data and analysis scripts used in the present study are available at the following link: https://osf.io/znvwb/. This study was not preregistered prior to data collection or analysis.

## Footnotes

1 Due to bioRxiv’s policy regarding human subjects, we cannot show an actual picture of the setup. See section 5 for supplemental videos of the task.

